# Modulation of mutant *Kras^G12D^*-driven lung tumorigenesis *in vivo* by gain or loss of PCDH7 function

**DOI:** 10.1101/343103

**Authors:** Xiaorong Zhou, Bret M. Evers, Mahesh S. Padanad, James A. Richardson, Emily Stein, Jingfei Zhu, Robert E. Hammer, Kathryn A. O’Donnell

## Abstract

PROTOCADHERIN 7 (PCDH7), a transmembrane receptor and member of the Cadherin superfamily, is frequently overexpressed in lung adenocarcinoma and is associated with poor clinical outcome. While PCDH7 was recently shown to promote transformation and facilitate brain metastasis in lung and breast cancers, decreased PCDH7 expression has also been documented in colorectal, gastric, and invasive bladder cancers. These data suggest context-dependent functions for PCDH7 in distinct tumor types. Given that PCDH7 is a potentially targetable molecule on the surface of cancer cells, further investigation of its role in tumorigenesis *in vivo* is needed to evaluate the therapeutic potential of its inhibition. Here we report the analysis of novel PCDH7 gain- and loss-of-function mouse models and provide compelling evidence that this cell-surface protein acts as a potent lung cancer driver. Employing a Cre-inducible transgenic allele, we demonstrated that enforced PCDH7 expression significantly accelerates *Kras^G12D^*-driven lung tumorigenesis and potentiates MAPK pathway activation. Furthermore, we performed *in vivo* somatic genome editing with CRISPR/Cas9 in *Kras^LSL-G12D^*; *Tp53^fl/fl^* (KP) mice to assess the consequences of PCDH7 loss of function. Inactivation of PCDH7 in KP mice significantly reduced lung tumor development, prolonged survival, and diminished phospho-activation of ERK1/2. Together, these findings establish a critical oncogenic function for PCDH7 *in vivo* and highlight the therapeutic potential of PCDH7 inhibition for lung cancer. Moreover, given recent reports of elevated or reduced PCDH7 in distinct tumor types, the new inducible transgenic model described here provides a robust experimental system for broadly elucidating the effects of PCDH7 overexpression *in vivo*.

**AUTHOR SUMMARY:** Lung cancer is the leading cause of cancer-associated deaths worldwide. PROTOCADHERIN 7 (PCDH7), cell surface protein and member of the Cadherin superfamily, is frequently overexpressed in lung adenocarcinomas and is associated with poor clinical outcome. Nevertheless, it has yet to be shown *in vivo* whether PCDH7 plays a role in the initiation and progression of lung cancer, and whether it represents an actionable therapeutic target. Here we demonstrate, using a novel transgenic mouse model, that PCDH7 overexpression accelerates *Kras^G12D^*-driven lung tumorigenesis. Furthermore, we validate PCDH7 as a therapeutic target by knocking it out using *in vivo* somatic genome editing in the *Kras^LSL-G12D^*; *Tp53^fl/fl^* (KP) model. Our results provide new insight into the mechanisms that drive lung cancer pathogenesis and, because targeting oncogenic cell-surface proteins with antibodies has proven to be a highly effective anti-cancer therapeutic strategy, establish a new target for cancer treatment. Moreover, given recent reports of elevated or reduced PCDH7 in distinct tumor types, the transgenic PCDH7 model described here provides a robust experimental system for elucidating the effects of PCDH7 overexpression in different *in vivo* settings. This model will also provide an ideal system for future testing of therapeutics directed at PCDH7.

## INTRODUCTION

Protocadherins (PCDHs) are transmembrane proteins and members of the Cadherin superfamily that play well-established roles in cell adhesion and regulation of downstream signaling pathways (1, 2). A growing body of evidence has demonstrated that PCDH expression is dysregulated in tumorigenesis (3-7). Both oncogenic and tumor suppressive roles have been assigned to PCDHs. However, the roles of individual PCDHs in cancer and the mechanisms through which their gain-of-function and loss-of-function drive tumorigenesis *in vivo* remain poorly understood. We previously reported that PROTOCADHERIN 7 (PCDH7) is frequently overexpressed in human non-small cell lung cancer (NSCLC) tumors (8). Moreover, high expression of PCDH7 was associated with poor clinical outcome of lung adenocarcinoma patients. While PCDH7 was recently shown to mediate brain metastasis in breast and lung cancers (9-12), decreased PCDH7 expression has been documented in colorectal, gastric, and invasive bladder cancers (13-15). Collectively, these data suggest context-dependent functions for PCDH7 in distinct tumor types. Because this molecule is a cell surface receptor that is potentially accessible to antibody-based therapies, rigorous *in vivo* studies are needed to interrogate PCDH7 function in cancer pathogenesis.

Lung cancer is the leading cause of cancer-associated deaths worldwide (16). Given the limited effectiveness of current treatments, there is a critical need to identify new therapeutic targets. PCDH7 transforms human bronchial epithelial cells (HBECs) and synergizes with *KRAS* to induce MAPK signaling and tumorigenesis in immunocompromised mice (8). One mechanism through which PCDH7 potentiates ERK signaling is by facilitating interaction of Protein Phosphatase 2A (PP2A) with its potent inhibitor, the SET oncoprotein, thereby suppressing PP2A activity (8). Depletion of PCDH7 suppressed ERK activation, sensitized NSCLC cells to MEK inhibitors, and reduced growth of lung cancer cells in xenograft assays. Nevertheless, all prior studies of PCDH7 function in cancer utilized established cell lines. Thus, investigation of the oncogenic activity of PCDH7 *in vivo* using autochthonous tumor models, which more accurately model multi-stage tumor progression and the role of the tumor microenvironment, is needed to dissect the role of this cell surface receptor in cancer pathogenesis and to evaluate the therapeutic potential of PCDH7 inhibition for non-small cell lung cancer.

In this study, we sought to establish the importance of PCDH7 in *Kras^G12D^*-driven lung cancer pathogenesis. To examine the effects of PCDH7 upregulation in lung tumor initiation and progression, we generated an inducible *PCDH7* transgenic mouse model, allowing the demonstration that hyperactivity of *PCDH7* promoted lung tumorigenesis and induced MAPK pathway activation in *Kras^LSL-G12D^* mutant mice (17). Additionally, to validate PCDH7 as a therapeutic target, we performed *in vivo* somatic gene editing in *Kras^LSL-G12D^*; *Tp53^fl/fl^* (KP) mice (18). Somatic depletion of PCDH7 in KP mice with CRISPR/Cas9 significantly reduced lung tumor development and prolonged survival, suggesting that targeting PCDH7 may benefit patients with lung adenocarcinoma. Interrogation of downstream signaling pathways revealed diminished phospho-ERK1/2 and phospho-RB in *PCDH7* knockout tumors. These findings demonstrate a key oncogenic role for PCDH7 *in vivo*, supporting the potential therapeutic efficacy of PCDH7 inhibition for patients with lung cancer.

## RESULTS

### PCDH7 accelerates lung tumorigenesis in a mouse model of *Kras^G12D^*-driven lung adenocarcinoma

To assess the oncogenic activity of PCDH7 *in vivo* and to investigate the mechanisms underlying PCDH7-mediated transformation, we generated a transgenic mouse model that allows precise control of PCDH7 expression through Cre-mediated recombination. We selected human PCDH7 isoform A, which is the predominant isoform expressed in human lung cancer cell lines from the Cancer Cell Line Encyclopedia and a panel of NSCLC cell lines (8, 19). The transgene is depicted in **Fig 1A**. A LoxP-Stop-LoxP (LSL) cassette allows control with Cre-recombinase. After removal of the LSL cassette, the CAG promoter drives expression of human PCDH7 and green fluorescent protein (GFP). To validate inducible expression of the transgene, *PCDH7^LSL^* mice were crossed to *CAG-Cre* mice that ubiquitously express Cre recombinase (**S1A Fig**). Cre-dependent transgene activation was confirmed in lung tissue using quantitative real-time PCR and western blot analysis (**S1A-C Fig**).

**Fig 1.**
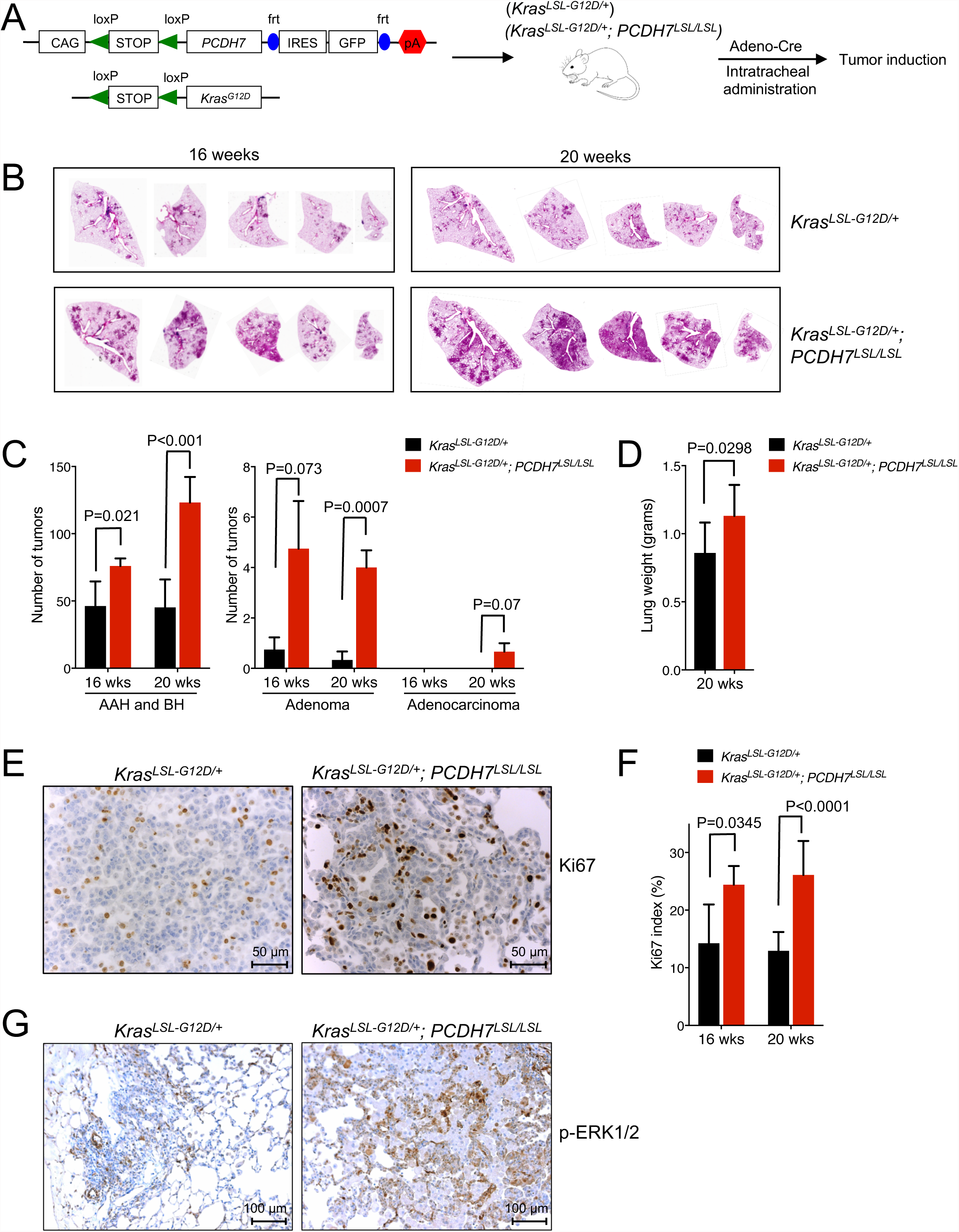
PCDH7 accelerates lung tumorigenesis in the *Kras^LSL-G12D^* model. (A) Schematic depicting the *PCDH7^LSL^* and *Kras ^LSL-G12D^* transgenes and overview of experimental design. To induce lung tumors, *Kras^LSL-G12D^; PCDH7^LSL/LSL^* or control *Kras^LSL-G12D^* mice were infected with Adeno-Cre through intratracheal administration. (B) H&E staining of lung lobes harvested at 16 and 20 weeks post-infection. Five independent lobes from one animal of each genotype are shown. (C) Tumor burden analysis at 16 and 20 weeks post-infection. n=5 mice/group. Tumors were counted and classified into atypical adenomatous hyperplasia (AAH)/bronchiolar hyperplasia (BH), adenomas, or adenocarcinomas. (D) Weight of lungs collected from *Kras^LSL-G12D^; PCDH7^LSL/LSL^* or *Kras^LSL-G12D^* mice at 20 weeks post-infection. (E) IHC staining of Ki67 at 20 weeks and (F) quantification of Ki67 index for lung sections harvested from *Kras^LSL-G12D^; PCDH7^LSL/LSL^* or control *Kras^LSL-G12D^* mice at 16 and 20 weeks post-infection. (G) IHC staining of pERK1/2 for lung sections from *Kras^LSL-G12D^; PCDH7^LSL/LSL^* or control *Kras^LSL-G12D^* mice at 16 weeks post-infection.

Replication-deficient viruses have been widely used to deliver Cre to the lung, thereby activating a mutant *Kras* allele to generate sporadic lung tumors in *Kras^LSL-G12D^* mice (17, 20-22). To determine whether enforced expression of PCDH7 is sufficient to accelerate *Kras^G12D^*-mediated lung tumorigenesis *in vivo*, we bred *PCDH7^LSL^* mice with *Kras^LSL-G12D^* mice and induced both transgenes via intratracheal administration of Adenovirus-Cre (23) (**Fig 1A**). Tumors were detectable in all lobes at 16 and 20 weeks post-infection (**Fig 1B**). At 16 weeks post-infection, the majority of tumors were early-stage and categorized as atypical adenomatous hyperplasia (AAH) or bronchiolar hyperplasia (BH) (17, 18, 23) (**Fig 1C** and **S1D Fig**). *Kras^LSL-G12D^*; *PCDH7^LSL/LSL^* mice exhibited a significant increase in AAH and BH tumors compared to *Kras^LSL-G12D^* mice. *Kras^LSL-G12D^*; *PCDH7^LSL/LSL^* mice also developed more adenomas than *Kras^LSL-G12D^* mice, indicating accelerated disease progression. At 20 weeks post-infection, tumor burden was significantly higher in *Kras^LSL-G12D^*; *PCDH7^LSL/LSL^* mice than in *Kras^LSL-G12D^* mice, as determined by quantifying tumor numbers at different stages (**Fig 1C**) and lung weight (**Fig 1D** and **S1E Fig**). No tumors were observed in *PCDH7^LSL/LSL^* mice after 16 months post-infection, suggesting that isolated expression of the transgene is not sufficient to initiate tumorigenesis by this time-point.

PCDH7 cooperates with oncogenic KRAS to promote ERK activation and cell proliferation in human bronchial epithelial cells (HBECs) (8). Consistent with these effects, tumors from *Kras^LSL-G12D^*; *PCDH7^LSL/LSL^* mice exhibited increased levels of Ki67 and pERK1/2 staining compared to tumors from *Kras^LSL-G12D^* mice (**Fig 1E-G**). Taken together, these data support a pro-tumorigenic role for PCDH7 in *Kras^G12D^-*mutant lung cancer and demonstrate that PCDH7 modulates MAPK pathway activity *in vivo*.

### *In vivo* inactivation of *Pcdh7* reduces lung tumor burden and prolongs survival of *Kras^LSL-^*^G12D^; *Tp53^fl/fl^* mice

Inhibition of PCDH7 reduced tumorigenesis of *KRAS* mutant NSCLC cells in xenograft assays (8). However, xenografts do not fully recapitulate all aspects of tumorigenesis, including contributions from the microenvironment and immune system. Therefore, examination of *in vivo* loss-of-function is critical to establish whether PCDH7 represents a potential therapeutic target in NSCLC. We took advantage of a recently described *in vivo* somatic genome editing approach to rapidly interrogate PCDH7 function in *Kras^LSL^*^G12D^;*Tp53^fl/fl^* (KP) mice (**Fig 2A**) (18). Interestingly, PCDH7 is upregulated in KP tumors relative normal lung, suggesting it may act as an oncogenic driver in this context (**S2A Fig**). CRISPR/Cas9-based gene editing efficiently produces loss-of-function mutations in this model that result in phenotypes that closely mirror those observed following traditional germline gene-targeting approaches.

**Fig 2.**
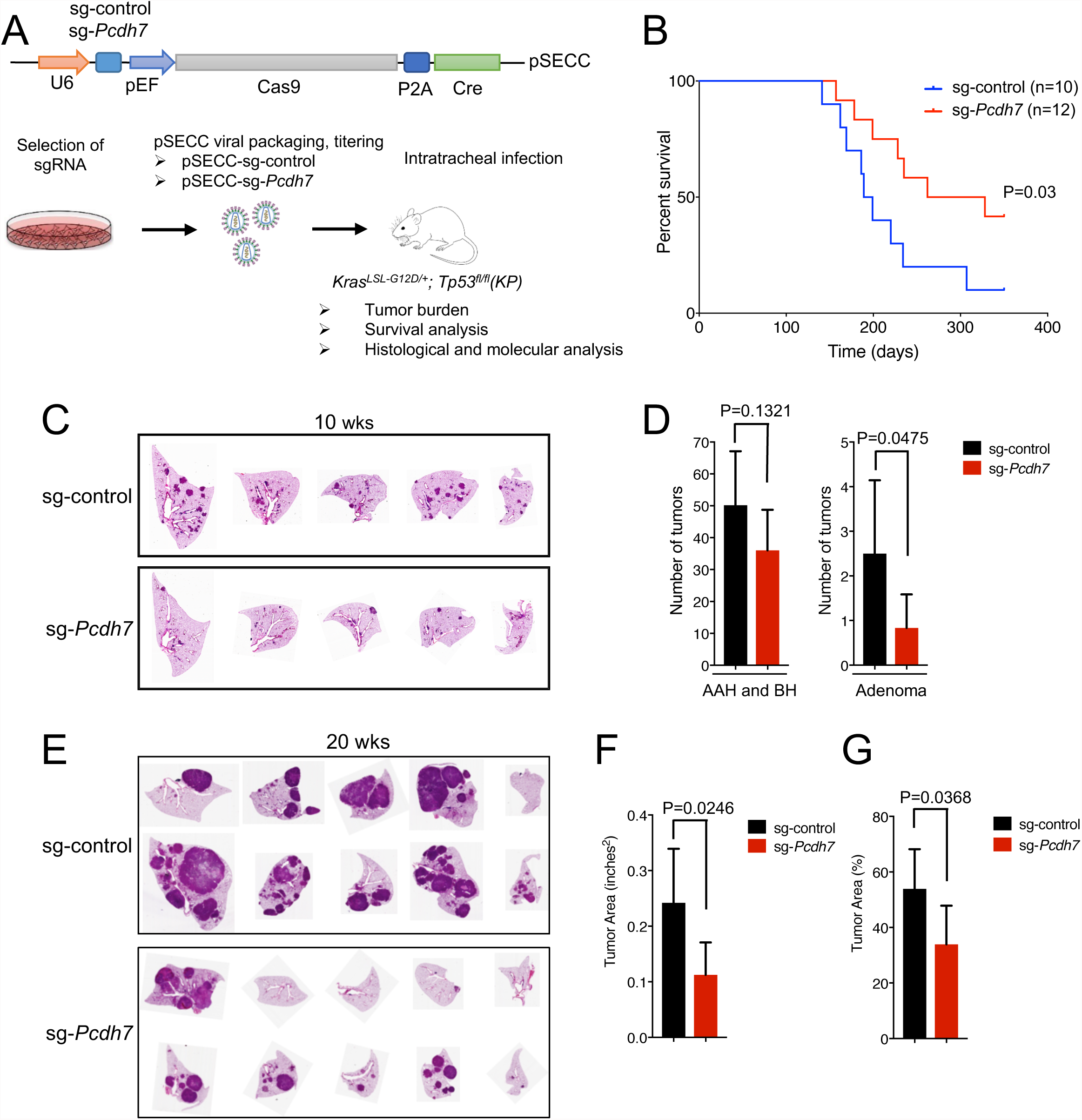
*In vivo* inactivation of *Pcdh7* reduces lung tumor burden and prolongs survival of *Kras^LSL-G12D^*; *Tp53^fl/fl^* mice. (A) Schematic of *in vivo* CRISPR/Cas9 editing. KP mice were infected with a sg-control or *sg-Pcdh7* through intratracheal administration of pSECC lentivirus expressing Cas9 and Cre. (B) Survival analysis of KP mice infected with sg-control or sg-*Pcdh7* lentivirus. (C) H&E staining of lung lobes harvested at 10 weeks post-infection. (D) Tumor burden analysis at 10 weeks post-infection. Tumors were counted and classified into AAH, BH, or adenomas based on histopathologic analysis. (E) H&E staining of lung lobes harvested at 20 weeks post-infection. (F) Total tumor area (inches^2^) at 20 weeks post-infection based on histopathologic analysis of lung sections. (G) Percent tumor area compared to total area of lung lobes at 20 weeks post-infection.

To identify an optimal *Pcdh7* single-guide (sgRNA) for *in vivo* studies, immortalized mouse embryonic fibroblasts were infected with a lentivirus expressing the Cas9 nuclease and a control sgRNA or one of five sgRNAs directed against murine *Pcdh7* (**Fig 2A**) (24). *Pcdh7* sgRNA-1 most dramatically diminished PCDH7 expression by western blot analysis (**S2B Fig**). CRISPR/Cas9-induced double strand DNA breaks were verified with the SURVEYOR assay (25) (**S2C Fig**), thus *Pcdh7* sgRNA-1 was selected for *in vivo* targeting.

*Kras^LSL-G12D^*; *Tp53^fl/fl^* (KP) mice were infected with pSECC-sg-control or pSECC-sg-*Pcdh7* lentiviruses through intratracheal administration, following established protocols (23). Because the lentivirus expresses Cas9 and Cre, the *Pcdh7* sgRNA generates mutations in the same lung cells that undergo activation of the *Kras^LSL-G12D^* allele and deletion of the*Tp53^fl/fl^* allele. Somatic mutation of *Pcdh7* prolonged survival of *Kras^LSL-G12D^*; *Tp53^fl/fl^* mice (**Fig 2B**). At an early time-point (ten weeks after infection), we observed fewer adenomas but not AAH or BH lesions in *Pcdh7-*targeted KP lungs (**Fig. 2C-D**), suggesting that depletion of PCDH7 impairs tumor progression. Moreover, at 20 weeks post-infection, a time-point associated with robust adenocarcinoma formation in KP mice, we documented significantly reduced tumor burden (**Fig 2E-G**).

Given recent studies documenting a role for PCDH7 in metastasis, we assessed invasive properties of tumors in these mice. We observed evidence of micro- and macro-metastases, lymphovascular invasion, and intravascular tumor spreading in control-treated *Kras^LSL-G12D^*; *Tp53^fl/fl^* mice (**S3 Fig**). sg*-Pcdh7*-treated mice exhibited a lower incidence of these malignant features and a higher incidence of relatively benign neoplastic lesions, including AAH and adenomas. Thus, *Pcdh7* depletion reduced tumor invasiveness *in vivo*.

To verify gene editing events, genomic DNA was harvested from *Pcdh7*-targeted tumors at 20 weeks after infection, mutations in the sgRNA-targeted region of *Pcdh7* were sequenced, and PCDH7 expression was measured by western blotting. All control tumors from *Kras^LSL-G12D^*; *Tp53^fl/fl^* mice expressed high levels of PCDH7 relative to normal lung tissue (**Fig 3A**). As expected, tumors isolated from sg-*Pcdh7*-targeted KP mice exhibited diminished PCDH7 expression to varying degrees across analyzed tumors (**Fig 3A** and **S4A Fig**). Accordingly, sequencing of the sg-*Pcdh7*-targeted region revealed homozygous or heterozygous deletions resulting in a frameshift in 8/9 tumors analyzed, with residual wild-type or in-frame alleles in tumors with higher PCDH7 expression (**S4B Fig**).

**Fig 3.**
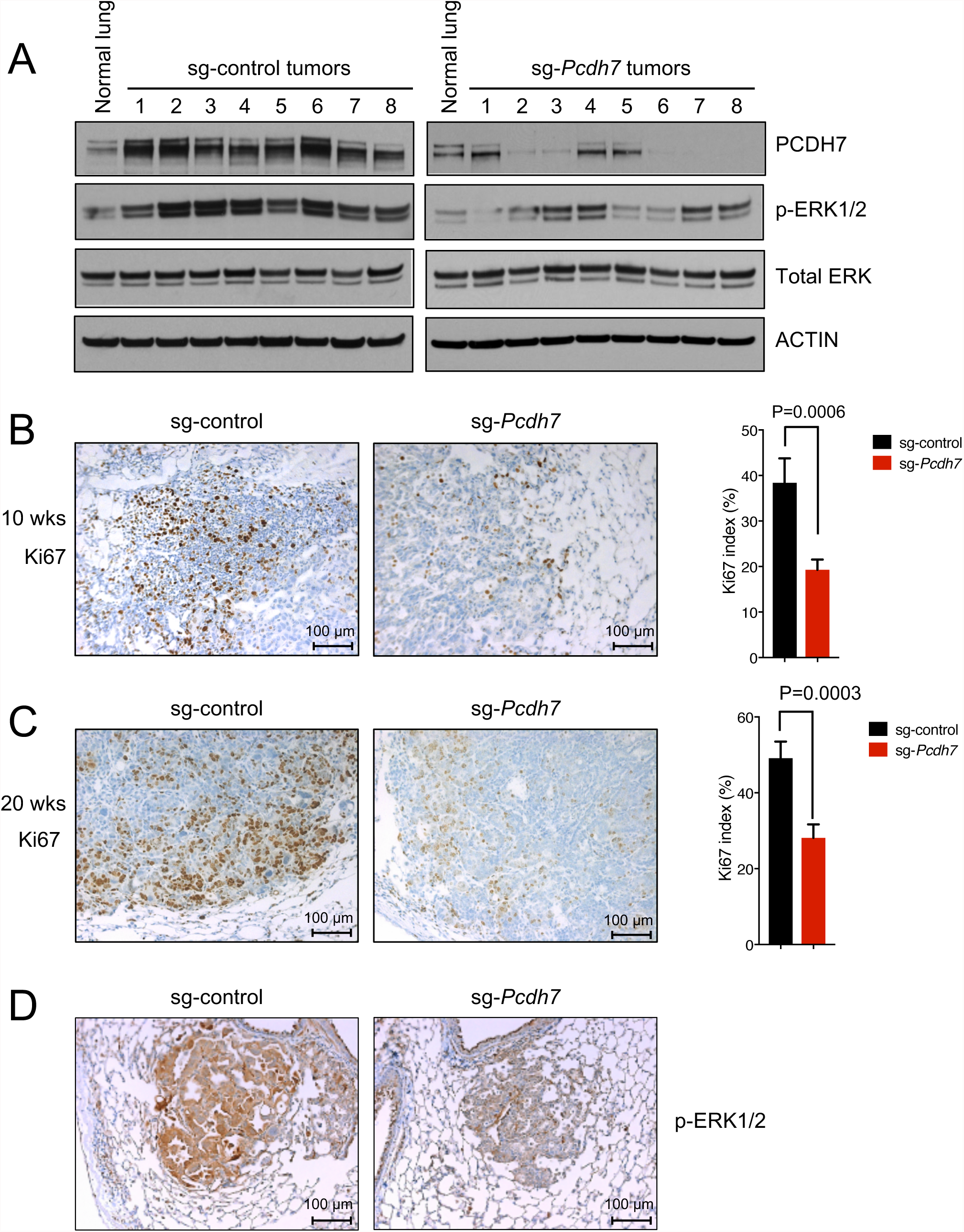
Somatic knockout of *Pcdh7* inhibits proliferation and reduces MAPK signaling in *Kras^LSL-G12D^*; *Tp53^fl/fl^* mice. (A) Western blot analysis of PCDH7, pERK1/2, and total ERK in normal lung tissues from uninfected KP mice, and lung tumors from KP mice infected with sg-control or *sg-Pcdh7* lentivirus (20 weeks post-infection). (B-C) *Left*, Ki67 immunohistochemistry (IHC) staining of lung sections harvested from KP mice infected with sg-control or sg-*Pcdh7* lentivirus at 10 weeks (B) or 20 weeks (C) post-infection. *Right*, Ki67 index = percent of cells with a Ki67 positive signal / total cell number in each field. n=2-4 animals per group, up to 9 independent fields were quantified for each animal. (D) pERK1/2 IHC staining of lung lobes from *Kras^LSL-G12D^; Tp53 fl/fl* mice infected with sg-control or sg-*Pcdh7* lentivirus at 10 weeks post-infection.

### Somatic knockout of *Pcdh7* inhibits proliferation and reduces MAPK signaling in *Kras*^*LSL*-G12D^; *Tp53^fl/fl^* mice

To examine the effects of PCDH7 depletion on downstream signaling *in vivo*, we harvested tumors from KP mice infected with either sg-control or sg-*Pcdh7* lentivirus and analyzed phospho-ERK1/2 and total ERK expression by western blotting (**Fig 3A**).

Although phospho-ERK1/2 was uniformly lower in sg-*Pcdh7*-targeted tumors compared to control tumors, some heterogeneity existed, consistent with additional genetic events affecting the MAPK pathway. Immunohistochemistry demonstrated that pERK1/2 and Ki67 staining were reduced in tumors from sg-*Pcdh7*-targeted mice compared to control KP tumors (**Fig 3B-D**), further supporting a role for PCDH7 in stimulating MAPK pathway activity.

### PCDH7 modulates expression of PP2A targets

PCDH7 overexpression in HBECs impact several important cancer-relevant pathways. One reported target of PP2A is pRB (26). Accordingly, the pRB pathway was previously identified as a significantly upregulated gene set in RNAseq analysis of HBEC-shp53-*PCDH7* cells (p=0; FDR q value = 2.35 × 10^−4^) (8). To determine whether PCDH7 regulates the pRB pathway in the context of mutant *KRAS^G12V^*, we generated HBEC-shp53 cells with enforced expression of mutant *KRAS^G12V^*, *PCDH7*, or both in combination. Western blot analysis revealed that while KRAS^G12V^ increased pERK and pRB signaling, PCDH7 cooperated with KRAS^G12V^ to further enhance both pERK and pRB signaling (**Fig 4A**). Furthermore, both pERK and pRB signaling were significantly reduced upon PCDH7 inhibition with siRNAs in shp53-*KRAS^G12V^-PCDH7* cells (**Fig 4B**).

**Fig 4.**
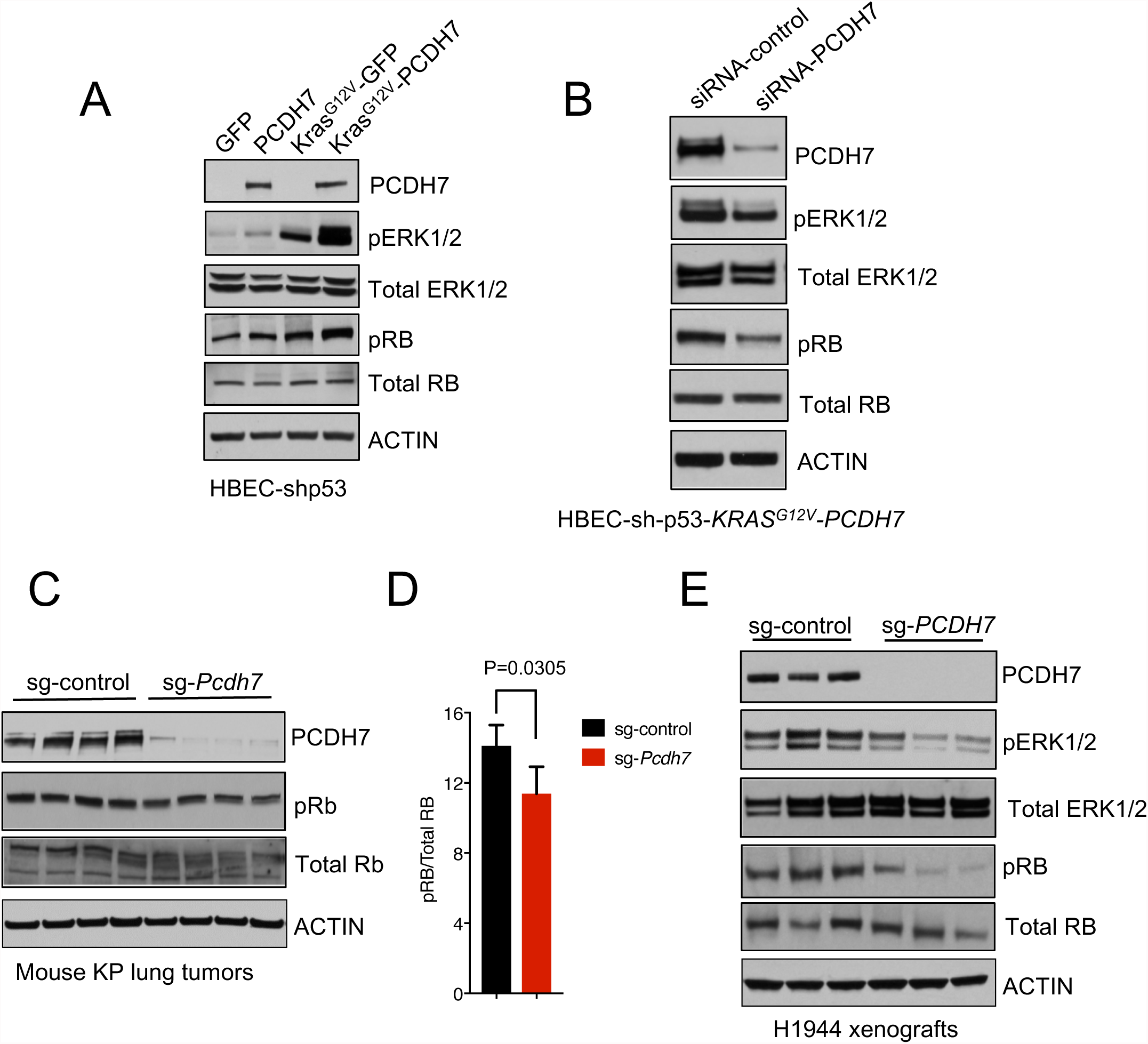
PCDH7 modulates expression of PP2A targets pERK1/2 and pRB. (A) Western blot analysis of PCDH7, pERK, total ERK, pRb and total RB levels in HBECs expressing *KRAS^G12V^*, *PCDH7*, or both. (B) PCDH7 inhibition with siRNAs in HBEC-shp53-*KRAS^G12V^-PCDH7* cells and its effects on pERK and pRb signaling, as indicated by western blot analysis. (C) Western blot analysis of PCDH7, pRB, and total RB expression in lung tumors harvested from KP mice with sg-control or sg-*Pcdh7* lentivirus at 20 weeks post-infection. (D) Quantification of pRB protein levels shown in (C). (E) Western blot analysis showing diminished pERK1/2 and pRB protein in human NSCLC H1944 xenografts with *PCDH7* sgRNA.

To determine the extent to which PCDH7 inhibition impacts the RB pathway *in vivo*, sg-control or sg-*Pcdh7*-treated tumors from KP mice were collected at 20 weeks post-infection, and pRB and total RB were examined by western blotting. A minor, but significant, decrease in pRB signaling was observed in PCDH7-depleted tumors compared to control tumors (**Fig 4C-D**). Finally, we extended these studies to human NSCLC xenografts. In *KRAS* mutant H1944 lung adenocarcinoma cells, CRISPR/Cas9-mediated depletion of PCDH7 suppressed the growth of xenografts in immunocompromised NOD/SCID IL2Rγ^null^ NSG mice (8). Western blotting demonstrated that PCDH7 depletion diminished both pRB and pERK signaling *in vivo* (**Fig 4E**). Collectively, these data show that PCDH7 enhances phosphoactivation of multiple PP2A targets including pERK1/2, and to a lesser extent pRB, in KRAS-mutant lung cancer cells.

## DISCUSSION

### PCDH7 is an actionable therapeutic target in lung adenocarcinoma

The inducible PCDH7 transgenic mouse described here provides a valuable model for the evaluation of PCDH7 function in various tumorigenic contexts. Given the prior reports of elevated or reduced PCDH7 in distinct tumor types (8, 10, 13-15), the *PCDH7^LSL^* model, when combined with various tissue-specific Cre driver lines, provides a robust experimental system for elucidating the effects of PCDH7 overexpression in different *in vivo* settings. Furthermore, this animal model represents an ideal system for future testing of therapeutics directed at PCDH7, including monoclonal antibodies.

We previously demonstrated that PCDH7 interacts with the PP2A phosphatase, and SET (a potent PP2A inhibitor), thereby inhibiting PP2A activity in HBECs and human NSCLC cells (8). PP2A loss of function results in aberrant phosphorylation of substrates in a variety of pathways linked to cancer, including the MAPK, RB, AKT, and JAK/STAT pathways (27-30). Our *in vivo* gain- and loss-of-function studies demonstrate that PCDH7 promotes ERK phosphoactivation, an event known to initiate lung tumorigenesis and promote the rapid progression of adenomas to more invasive adenocarcinomas (31). Importantly, the magnitude and duration of ERK signaling are tightly controlled by regulators that provide negative feedback, including PP2A (32, 33). Moreover, the consequences of PCDH7-mediated PP2A inhibition likely extend beyond MAPK signaling. Indeed, we show here that PCDH7 modulates pRB levels in human NSCLC xenografts and, to a lesser extent, in KP mouse tumors. Taken together, our data establish PCDH7 as a cooperative oncogenic driver in *KRAS*-mutant lung cancer that functions to enhance pro-tumorigenic signaling in cancer cells.

PCDH7 was recently shown to promote metastasis of lung and breast cancer cells (10, 34), further supporting the importance of this protein in multiple aspects of tumor biology. Consistent with these observations, our studies revealed that *Pcdh7* depletion in KP mice reduced tumor invasiveness *in vivo*. Overall, the results reported here establish PCDH7 as an oncogenic driver of lung tumor initiation and progression *in vivo*, setting the stage for future efforts to target this cell-surface protein with novel therapeutic strategies in lung adenocarcinoma.

## MATERIALS AND METHODS

### Constructs

Human PCDH7 isoform-A (NM_002589.2) cDNA was cloned into the CTV vector (Addgene #15912). The resulting CTV-PCDH7 was used to generate transgenic mice. Depletion of PCDH7 *in vitro* was performed by using Lenti-CRISPR-V2 (Addgene #52961) to introduce Cas9 and sgRNA directed against *Pcdh7*. Depletion of mouse *Pcdh7 in vivo* was performed using pSECC (Addgene #60820). The sgRNAs are as follows: human PCDH7, CGACGTCCGCATCGGCAACG; Mouse *Pcdh7*, GAGGATGCGGACCACGGGAT; human non-specific control, CGCTTCCGCGGCCCGTTCAA; mouse non-specific control, GCGAGGTATTCGGCTCCGCG. Human *Kras^G12V^* cDNA was cloned into pLenti CMV Hygro DEST (Addgene #17454) for ectopic expression of *Kras^G12V^* in HBECs. The human PCDH7 isoform-A cDNA was cloned into pLX303 lentivirus vector (Addgene #25897) for PCDH7 overexpression *in vitro*.

### Mice

*Generation and genotyping of PCDH7 transgenic mice -* PCDH7 transgenic mice were generated by UTSW transgenic core facility, through microinjection of the linearized CTV-PCDH7 vectors into the pronuclei of fertilized eggs (C57BL/6J strain). For genotyping of PCDH7 transgenic mice, genomic DNA was isolated from tail clippings using the Gentra Puregene Tissue Kit (Qiagen) according to the manufacturer’s protocol. PCR was performed using 10–100 ng of genomic DNA as template and the following transgene specific primers: forward: 5′-TCCCCAGTCACCAACTGCAGGAAAAAAACACCAG-3′, reverse: 5′-GAATAGGAACTTCGGTACCGAATTGATCGCG-3′ (amplicon = 299 bp). Thermal cycling conditions using GoTaq Green Master Mix (Promega) were as follows: initial denaturation at 95 °C for 5 min, followed by 95°C for 30 s, 60°C for 30 s, 72°C for 30 s, in a total of 35 cycles, followed by a final extension at 72°C for 5 min. The samples were stored at 4°C until separated on a 1.2% (wt/vol) agarose gel.

B6.129S4-Krastm4Tyj/J mice, also known as *Kras^LSL-G12D^* mice were purchased from The Jackson Laboratory (008179). *Kras^LSL-G12D^; PCDH7^LSL/LSL^* (maintained on a B6 genetic background) were established by breeding *Kras^LSL-G12D^* with *PCDH7^LSL^* mice. *Kras^LSL-G12D;^ Tp53^fl/fl^* mice (maintained on a FVB/B6 mixed genetic background) were provided by James Kim (UTSW). Immunocompromised mice NOD.Cg-Prkdcscid Il2rgtm1Wjl/SzJ (NSG mice, 005557) were purchased from The Jackson Laboratories. CAG-Cre mice (35) were provided by Eric Olson (UTSW).

### Ethics Statement

Mice were monitored closely throughout all experimental protocols to minimize discomfort, distress, or pain. Signs of pain and distress include disheveled fur, decreased feeding, significant weight loss (>20% body mass), limited movement, or abnormal gait. If any of these signs were detected, the animal was removed from the study immediately and euthanized. All sacrificed animals were euthanized with CO2. The animals were placed in a clear chamber and 100% CO2 was introduced. Animals were left in the container until clinical death ensured. To ensure death prior to disposal, cervical dislocation was performed while the animal was still under CO2 narcosis. All methods were performed in accordance with the recommendations of the Panel on Euthanasia of the American Veterinary Medical Association and protocols approved by the UT Southwestern Institutional Animal Care and Use Committee (protocol # 2017-102112).

### Tissue collection and immunohistochemistry

Mice were euthanized by intraperitoneal administration of an overdose of Avertin at the time points indicated. Lungs were inflated and perfused through the trachea with 4% paraformaldehyde (PFA), fixed overnight, transferred to 50% ethanol and subsequently embedded in paraffin. Sections were cut and stained with H&E by the UTSW Histology Core, which also provided assistance with the histopathological examination. For immunohistochemistry (IHC) staining, slides were de-paraffinized and treated with Antigen Unmasking Solution (Vector laboratory, H-3300), blocked with BLOXALL™ Blocking Solution (Vector lab, SP-6000), and then incubated with anti-Ki67 (CST #9027, 1:400) or anti-pERK (CST #9101S, 1:200) at 4°C overnight. After washing extensively, slides were incubated with SignalStain^®^ Boost Detection Reagent from Cell Signaling Technology (Rabbit #8114) at room temperature (RT) for 30 minutes. The signal was developed with ImmPACT™ DAB Substrate (Vector lab, SK-4105), and sections were counterstained with hematoxylin (Vector lab, H-3404), and mounted with VectaMount Mounting Reagent (Vector lab, H-5000). All pictures were obtained using a Zeiss microscope (Observer.Z1) with an Axiocam-MRC camera.

*Ki67 index* – Ki67-positive staining was distinguished by counting brown nuclei and hematoxylin (blue) counterstain. The Ki67 index for each mouse was calculated as follows: Percent positive cells = number of positive nuclei/total cell nuclei × 100, as described previously (36). 2-4 animals per group were analyzed, and up to 9 independent fields were quantified for each animal.

### Tumor burden analysis

For tumor quantification (Figure 1*C* and 2*D*), the total tumor number for each animal was determined by analyzing H&E staining and categorizing tumors into different stages. 5 lobes per animal were quantified, and 5-6 animals were analyzed per group. To analyze tumor burden of CRISPR/Cas9-targeted mice (Figure 2*F* and 2*G*), the total area of the tumor compared to the area of the whole lung for each mouse was obtained by analyzing H&E staining (5 lobes per animal) using the NIH Image J Program. The average tumor area (inches^2^) and percent of tumor vs. total lung area are shown (n=5-7 animals per group).

### Cell culture

Immortalized normal human bronchial epithelia cells (HBECs) with stable knockdown of *TP53* with shRNA (HBEC-shp53) were provided by John Minna at UTSW (37-39). Ectopic expression of *KRAS^G12V^* in HBEC-shp53 was performed with lentiviral infection followed by selection with hygromycin B for 7 days. HBEC-shp53-*PCDH7* or HBEC-shp53-*KRAS^G12V^-PCDH7* cells were established with the pLX303-PCDH7 lentivirus followed by selection with blasticidin for 10 days. All HBECs were cultured in keratinocyte serum-free medium (KSFM; Life Technologies Inc.) containing 50 μg/mL of bovine pituitary extract (BPE; Life Technologies, Inc.) and 5 ng/mL of EGF (Life Technologies, Inc.). Human lung adenocarcinoma H1944 cells were cultured in RPMI-1640 media supplemented with 10% fetal bovine serum (Life Technologies, Inc.), 100 units/ml of penicillin and streptomycin (Life Technologies, Inc.). PCDH7 depletion in H1944 cells was achieved using Lenti-CRISPRv2 followed by selection with puromycin for 7 days (1 μg/mL). All cell lines used in this study were cultured in 5% CO2 at 37C and tested negative for mycoplasma contamination.

### Virus preparation, titration and intratracheal administration

Cre-expressing adenovirus (Adeno-Cre) was purchased from Viral Vector Core Facility (University of Iowa). The *PCDH7^LSL^* and *Kras^LSL-G12D^* transgenes were induced with intratracheal administration of Adenovirus-Cre (Adeno-Cre, 2×10^7^ IFU/mouse) (23).

*Pcdh7* sgRNA-1 (sg-*Pcdh7*) or a non-specific control sgRNA (sg-control) were cloned into pSECC lentiviral vectors, packaged, and titered with Green-Go cells as previously described (18). Lentiviruses were produced by co-transfection of 293T cells with lentiviral backbone constructs and packaging vectors (psPAX2 and pMD2.G) using Lipofectamine 3000 (Invitrogen). Supernatants were collected 72 hours posttransfection, concentrated by ultracentrifugation at 25,000 RPM for 120 minutes and resuspended in an appropriate volume of Opti-MEM (Gibco). Green-Go cells, which were generated by transducing retrovirus containing an inverted GFP (flanked by two sets of incompatible loxP sites), were provided by Tyler Jacks (Massachusetts Institute of Technology), and maintained in DMEM supplemented with 10% Fetal Bovine Serum and gentamicin (18). Upon infection with pSECC lentivirus, Green-Go cells become GFP^+^, allowing for titering by fluorescence-activated cell sorting (FACS). Briefly, 2×10^4^ Green-Go cells were seeded into each well of 24-well plates and incubated overnight.

The next day, cells were infected with 20ul, 10ul, 5ul or 2ul pSECC lentivirus. At 48 hours post-infection, cells were collected and GFP^+^ cells were quantified by FACS. The titer of pSECC lentivirus is designated as the average number of GFP^+^ cells in the four groups. Intratracheal administration of pSECC lentivirus was performed according to established protocols (23). 4×10^4^ virus/mouse was administered for survival analysis and 2×10^5^ virus/mouse was administered for tumor burden analysis at 10 and 20 weeks post-infection (23).

### siRNA knockdown

PCDH7 siRNA (Accell™ SMARTpool) or non-specific control siRNA were purchased from Dharmacon. HBECs were incubated in Accell Delivery Media (Dharmacon) in 6 well-plates overnight to reach 50% confluence, and siRNAs diluted in siRNA Buffer (Dharmacon) were added at final concentration of 1μM. 72 hours after transfection, cells were collected, and gene expression was assessed by western blot.

### Western blot analysis

Protein lysates were prepared with RIPA Lysis and Extraction Buffer (ThermoFisher, 89900) including Halt™ Protease Inhibitor Cocktail, EDTA-free diluted 1:100. Protein concentrations were determined by BCA assay (Thermo, 23228 and 23224). 25-30µg of each protein lysate was loaded into each well of a Bolt™ 4-12% Bis-Tris Plus Gels (Life Technologies, NW04120BOX), electrophoresed, and transferred to nitrocellulose using iBlot2 Western Blotting System (Life Technologies). The primary antibodies (1:1000 dilution) for western blot are as follows: PCDH7 (Abcam, ab139274), pERK (Cell Signaling Techonolgy, CST # 9101S), total ERK (CST #4695), pRb (Ser807/811, CST #8516), total Rb (CST #9309) and β-ACTIN (CST #4970). Horseradish peroxidase (HRP)–conjugated anti-rabbit or anti-mouse secondary antibodies (1:5,000-20,000 dilution; BioRad) were used and signals developed with the SuperSignal™ West Dura Extended Duration Substrate (Thermo Fisher 34075), according to the manufacturer’s instructions.

### Verification of gene editing

To detect CRISPR-induced indels, tumors from a sg-*Pcdh7* mouse (20 weeks after infection) were dissected and surrounding lung tissues carefully removed. Tumor tissues were homogenized, and genomic DNA (gDNA) was isolated using the Gentra Puregene Tissue Kit (Qiagen). To amplify the *Pcdh7* sgRNA-targeted region, PCR was performed with tumor gDNA using PrimeSTAR HS DNA Polymerase (Clontech). The products were gel purified and cloned into the Zero Blunt TOPO sequencing vector (Invitrogen). At least eight colonies for each tumor were sequenced.

### Statistical Analysis

All statistical analyses were performed using GraphPad Prism for Windows. Quantitative variables were analyzed by Student’s t-test, Fisher’s exact test, or Chi-squared test. All statistical analyses were two-sided, and p<0.05 was considered statistically significant.

## Acknowledgements

We thank John Shelton in the University of Texas Southwestern Histology Core for assistance with histology, John Minna and Michael Peyton for sharing cell lines, and Tyler Jacks, David McFadden, James Kim, and Eric Olson for sharing reagents and mice. We also thank Joshua Mendell and members of the O’Donnell laboratory for critical reading of the manuscript. This work was supported by the NCI (R01 CA207763 to K.A.O.), the Sidney Kimmel Foundation (SKF-15-067 to K.A.O), the Cancer Prevention Research Institute of Texas (CPRIT, R1101 and RP150676 to K.A.O.), The Welch Foundation (I-1881 to K.A.O.), the LUNGevity Foundation (2015-03 to K.A.O.), and a SPORE in Lung Cancer CDA (P50CA70907-17). K.A.O. is a CPRIT Scholar in Cancer Research and a Kimmel Scholar. X. Zhou was supported by the Lung Cancer Research Foundation (LCRF 2015) and the National Natural Science Foundation of China (NSFC 81571527, 81771681).

## Author contributions

Conception and design of the work: K.A.O., X.Z., R.E.H.; Acquisition of data: X.Z., M.S.P., E.S., J.Z.; Analysis and interpretation of data: K.A.O., X.Z., B.M.E., J.A.R., R.E.H., M.S.P.; Writing and revision of manuscript: K.A.O., X.Z., B.M.E., J.A.R., R.E.H., M.S.P.

## Supplementary Figure Legends

**Fig S1.**
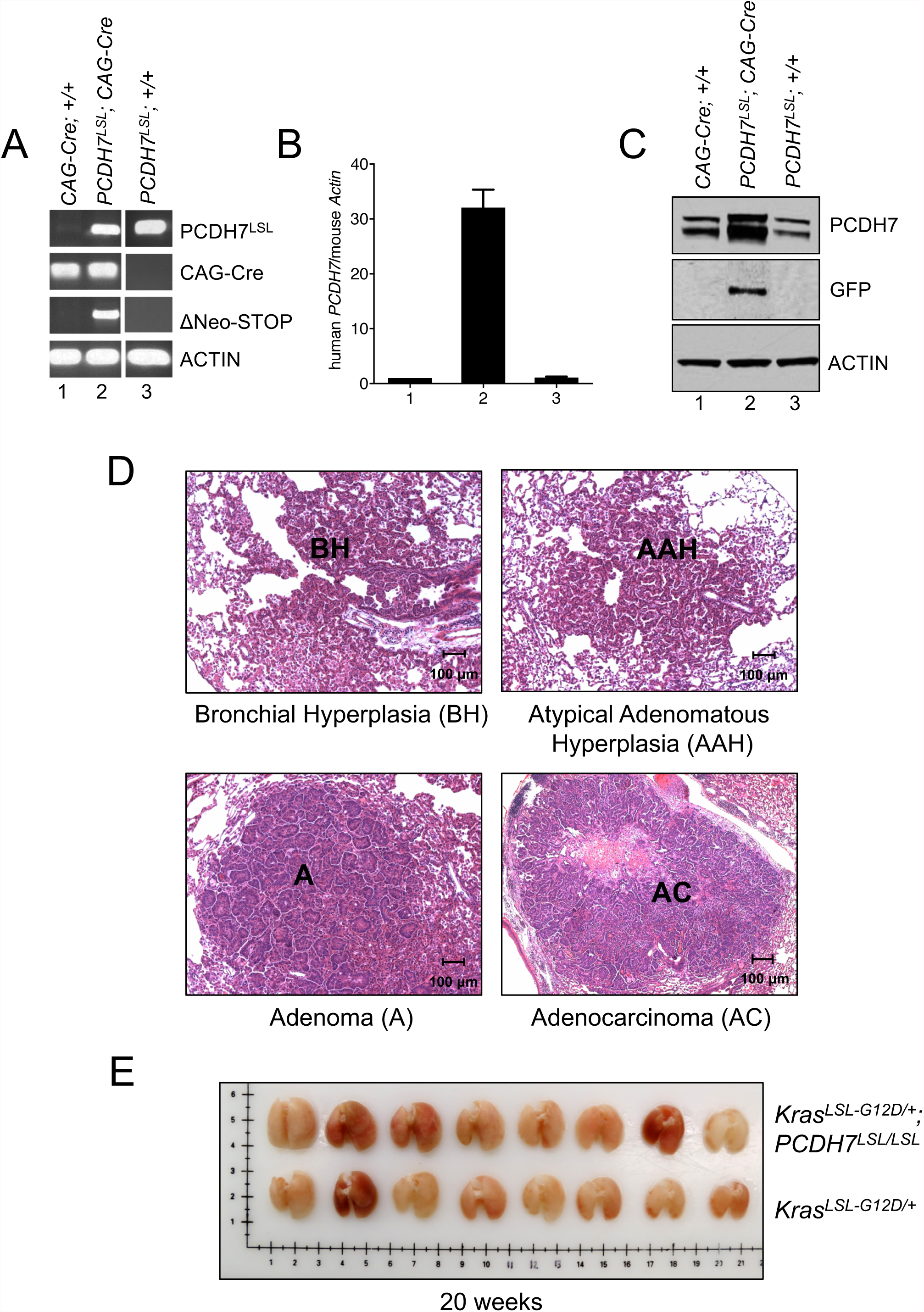
Generation of *PCDH7^LSL^* mice and analysis of lung tumors *in vivo*. (A) PCDH7 transgenic mice (*PCDH7^LSL^*) were crossed with CAG-Cre mice that ubiquitously express Cre under a CAG promoter. Genotyping PCR assay verifying Cre-mediated recombination, as indicated by deletion of a Neo-STOP cassette flanked by two LoxP sites. (B) qRT-PCR measurements of *PCDH7* transgene expression normalized to *Actin*. Error bars represent standard deviation of technical triplicates. (C) Western blot confirming transgenic PCDH7 and GFP expression in lung tissues. (D) Representative lung H&E images showing AAH, BH, adenomas, and adenocarcinomas in *Kras^LSL-G12D^; PCDH7^LSL/LSL^* mice (20 weeks post-infection). (E) Whole lung tissue isolated from *Kras^LSL-G12D^; PCDH7^LSL/LSL^* and control *Kras^LSL-G12D^* mice at 20 weeks post-infection.

**Fig S2.**
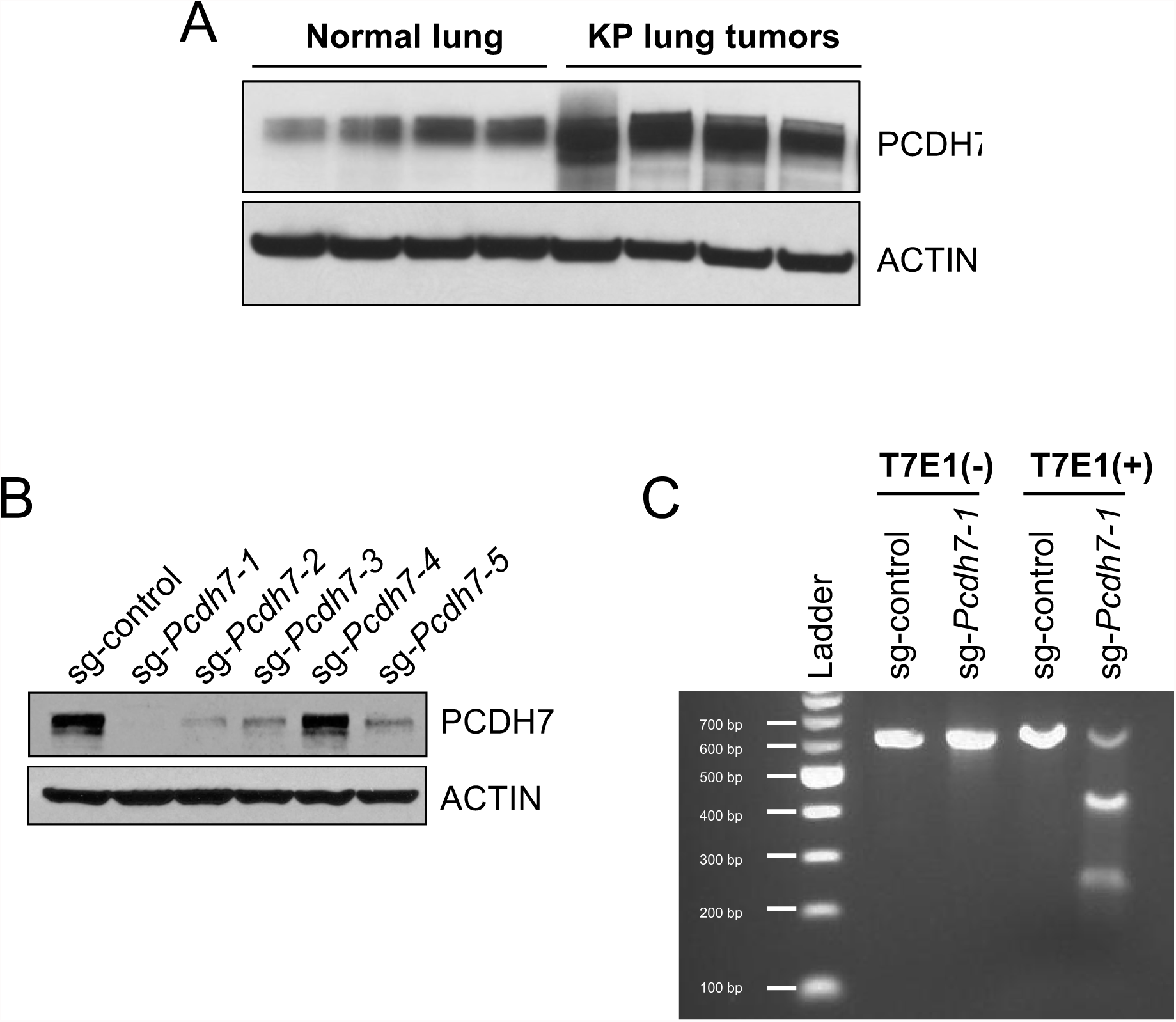
Validation of CRISPR/Cas9-editing of *Pcdh7.* (A) Western blot analysis of PCDH7 expression in normal mouse lung and lung tumors from *Kras^LSL-G12D^*; *Tp53^fl/fl^* mice. PCDH7 is upregulated 4-fold in four independent KP tumors. (B) Immortalized mouse embryonic fibroblasts (MEFs) infected with lentivirus with a control sgRNA or five individual *Pcdh7*-targeting sgRNAs. Western blot analysis of PCDH7 expression in puromycin-resistant cells. (C) Confirmation of genome modifications following gene editing of *Pcdh7* using the SURVEYOR assay. Green Go cells were infected with pSECC containing sg-control or sg-*Pcdh7-1*. GFP+ cells were FACS sorted after 7 days.

**Fig S3.**
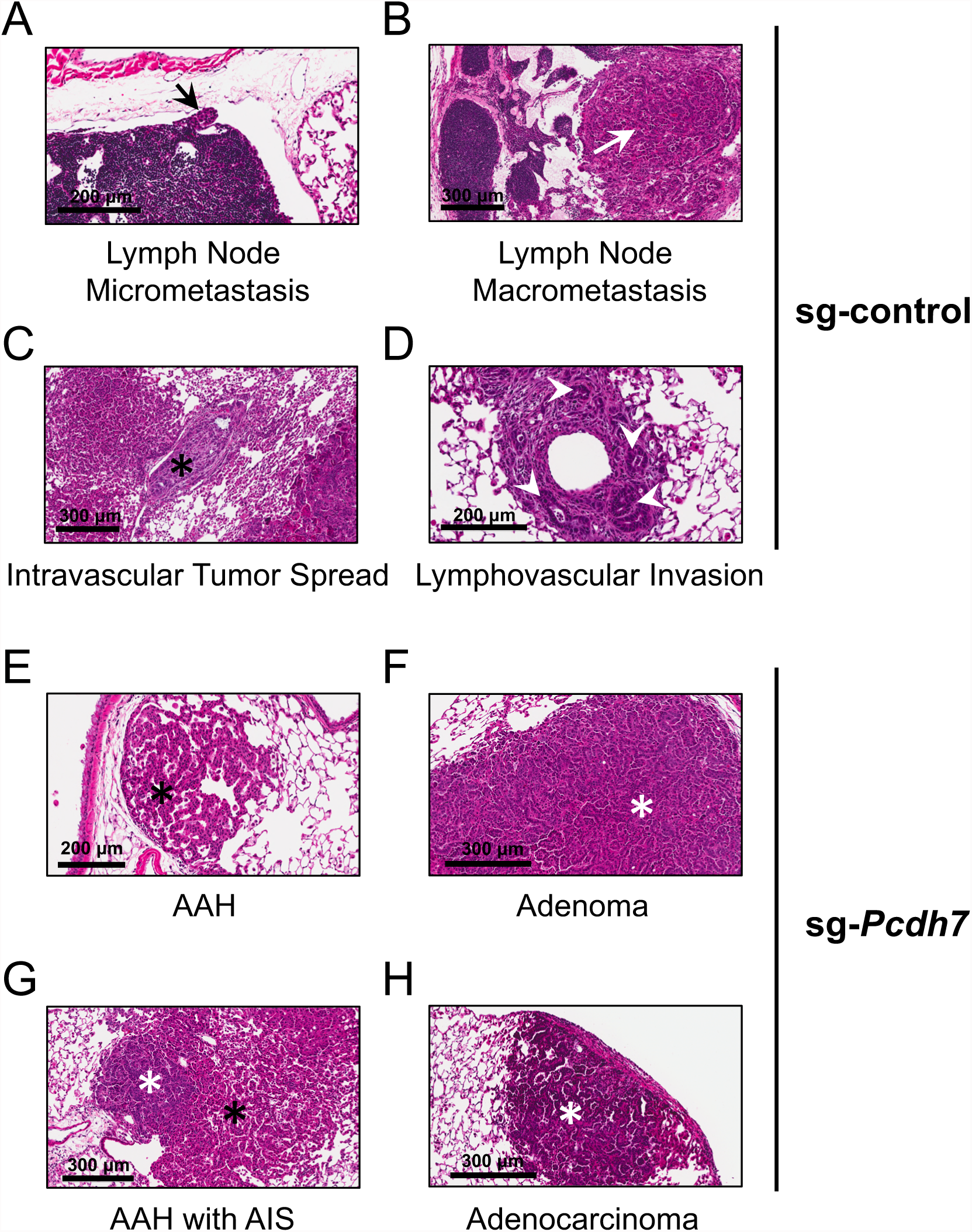
Diminished invasiveness in *Pcdh7* depleted KP tumors. Histology of lung lesions from sg-control and sg-*Pcdh7* mice. (A-D) H&E-stained lung sections from *sg*control mice showed features of advanced malignant tumor behavior, including both lymph node micrometastases (A, black arrow) and macrometastases (B, white arrow), intravascular adenocarcinoma spread (C, black asterisk), and lymphovascular invasion by adenocarcinoma (D, white arrowheads). (E-H) Conversely, H&E lung sections from sg*-Pcdh7* mice showed a lower incidence of these malignant features and a higher incidence of relatively benign neoplastic lesions, such as atypical adenomatous hyperplasia (AAH) (E and G, black asterisks) and adenomas (F, white asterisk). Though present at decreased frequencies, both adenocarcinoma in situ (AIS) (G, white asterisk) and frank adenocarcinoma (H, white asterisk) were also identified in sg-*Pcdh7* mice.

**Fig S4.**
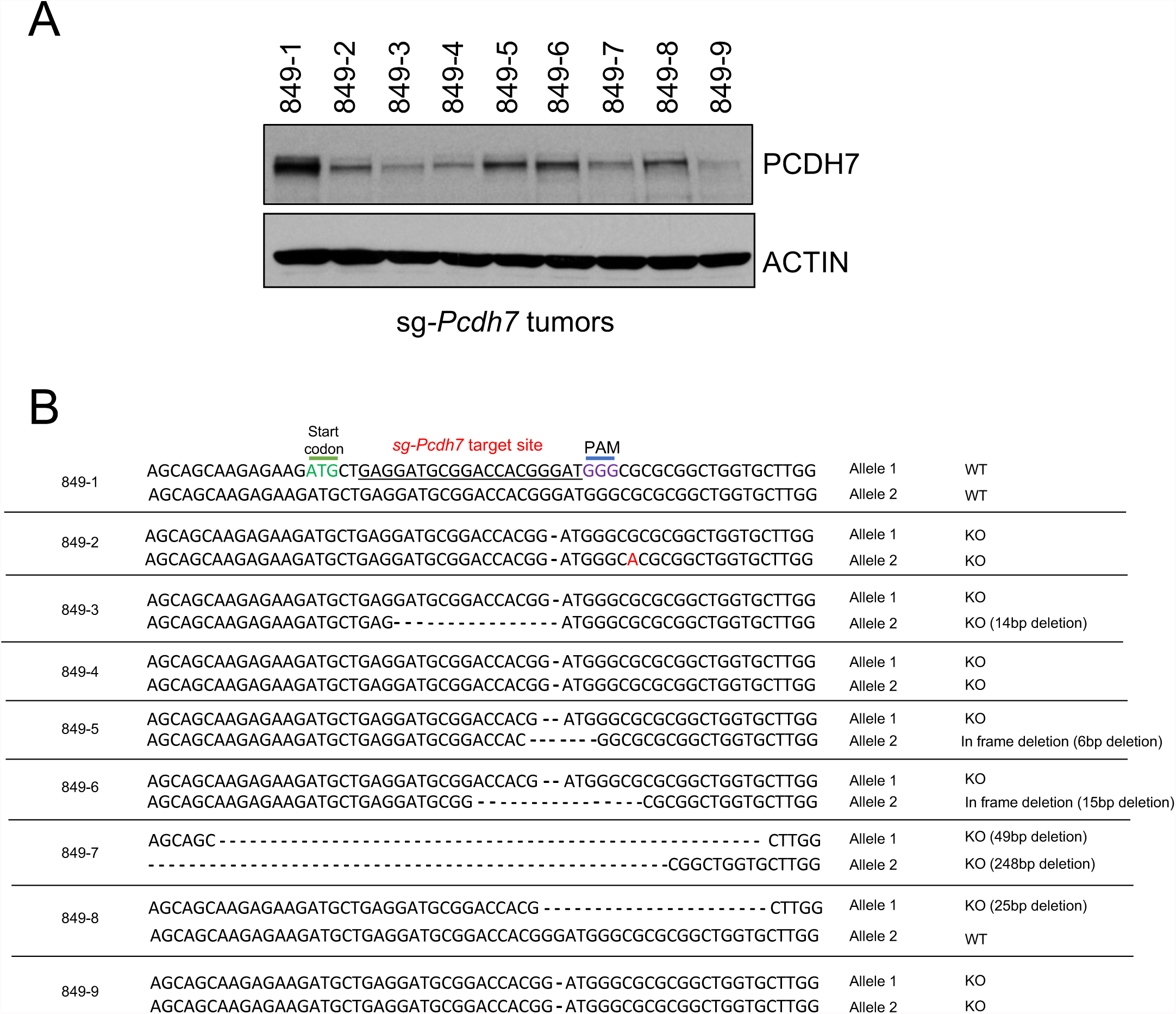
Sequences of gene-edited alleles in KP tumors. (A) Western blot analysis of PCDH7 protein in nine individual tumors from sg-*Pcdh7* targeted KP mice. (B) Mutations in the sg-*Pcdh7* targeted region were confirmed by cloning and sequencing.

